# Methylglyoxal-induced glycation stress promotes aortic stiffening: Putative mechanistic roles of oxidative stress and cellular senescence

**DOI:** 10.1101/2025.01.06.631561

**Authors:** Parminder Singh, Ravinandan Venkatasubramanian, Sophia A. Mahoney, Mary A. Darrah, Katelyn R. Ludwig, Alice Zhang, Kiyomi Kaneshiro, Lizbeth Enriquez Najera, Lauren Wimer, Muniesh M. Shanmugam, Edgard Morazan, Marrisa Trujillo, James Galligan, Richmond Sarpong, Douglas R. Seals, Pankaj Kapahi, Zachary S. Clayton

**Affiliations:** Buck Institute for Research on Aging, Novato, CA; University of Colorado Boulder, Boulder, CO; College of Chemistry, University of Berkely, Berkely, CA; R. Ken Coit College of Pharmacy, University of Arizona; University of Colorado Anschutz Medical Campus, Aurora, CO

## Abstract

**Background:** Here, we assessed the role of the advanced glycation end-product (AGE) precursor methylglyoxal (MGO) and its non-crosslinking AGE MGO-derived hydroimidazolone (MGH)-1 in aortic stiffening and explored the potential of a glycation stress-lowering compound (Gly-Low) to mitigate these effects.

**Methods:** Young (3–6 month) C57BL/6 mice were supplemented with MGO (in water) and Gly-Low (in chow). Aortic stiffness was assessed in vivo via pulse wave velocity (PWV) and ex vivo through elastic modulus. Putative mechanisms underlying MGO- and MGH-1-induced aortic stiffening were explored using complementary experimental approaches in aortic tissue and cultured human aortic endothelial cells (HAECs). Moreover, aortic stiffness was assessed in old (24 month) mice after consumption of Gly-Low-enriched chow.

**Results:** MGO-induced glycation stress increased PWV in young mice by 21% (P<0.05 vs. control), which was prevented with Gly-Low (P=0.93 vs. control). Ex vivo, MGO increased aortic elastic modulus 2-fold (P<0.05), superoxide production by ∼40% (P<0.05), and MGH-1 expression by 50% (P<0.05), which were all mitigated by Gly-Low. Chronic MGO exposure elevated biomarkers of cellular senescence in HAECs, comparable to a known senescence inducer Doxorubicin, an effect partially blocked by Gly-Low. Moreover, elevated aortic elastic modulus induced by Doxorubicin (P<0.05 vs. control) was prevented with Gly-Low (P=0.71 vs. control). Aortic RNA sequencing implicated preservation of endogenous cellular detoxification pathways with Gly-Low following exposure to MGH-1. Old mice supplemented with Gly-Low had lower PWV (P<0.05) relative to old control mice.

**Conclusions:** MGO-induced glycation stress contributes to aortic stiffening and glycation stress lowering compounds hold promise for mitigating these effects.

**What Is New?:** This study provides the first comprehensive line of evidence that methylglyoxal (MGO)-induced glycation stress directly contributes to aortic stiffening and does so through mechanisms involving oxidative stress and cellular senescence. Using complementary *in vivo*, *ex vivo*, and *in vitro* experimental models, we establish that MGO-mediated glycation stress independently induces aortic stiffening. Furthermore, we demonstrate that the glycation-lowering compound, Gly-Low, mitigates MGO-induced aortic stiffening by mitigating excessive oxidative stress and cellular senescence, and can lower aortic stiffness in old mice. Mechanistically, activation of the detoxification enzyme, glyoxalase-1 (Glo-1), is a novel pathway by which Gly-Low mediates its therapeutic effects on aortic stiffening. Lastly, we show that Gly-Low holds promise for lowering aortic stiffness in old age.

**What Is Relevant?:** Aortic stiffening is a major risk factor for cardiovascular diseases (CVD) and a significant predictor of CV-related morbidity and mortality. Yet, the underlying mechanisms driving this process remain incompletely understood. This study identifies MGO-derived glycation stress as a critical and modifiable factor contributing to aortic stiffening through pathways involving excessive oxidative stress and cellular senescence. By establishing the efficacy of Gly-Low in mitigating these effects, our findings underscore the importance of targeting glycation stress in the context of aging, and likely in other settings of glycation stress, to improve arterial health and reduce CVD risk.

**Clinical/Pathophysiological Implications:** These findings have significant clinical implications, as they demonstrate that glycation stress is a viable and modifiable therapeutic target for the prevention and treatment of aortic stiffening. Gly-Low offers a promising therapeutic approach to ameliorate glycation stress- and age-related aortic stiffening, by directly targeting excess glycation stress, oxidative stress, and cellular senescence. Additionally, the involvement of the Glo-1 detoxification pathway suggests a specific molecular target for future interventions aimed at improving arterial health and mitigating the progression of CVD.

## Introduction

Cardiovascular diseases (CVD) remain the leading cause of mortality worldwide, with elevated aortic stiffness being a significant predictor of CV-related morbidity and mortality^1^. Advancing age is the primary non-modifiable risk factor for the development of CVD. A key event mediating the increase in CVD risk with aging is stiffening of the large elastic arteries (primarily the aorta)^2^.

Age-related aortic stiffening, as demonstrated by increased aortic pulse wave velocity (PWV), occurs in part via increased accumulation of advanced glycation end products (AGEs), which results from non-enzymatic glycation of proteins and lipids^3,4^. Despite the biomedical importance AGEs, the underlying molecular mechanisms contributing to AGEs-mediated aortic stiffening with aging are not fully understood.

Methylglyoxal (MGO), a reactive α-dicarbonyl compound, is a key potent precursor of AGEs and is significantly elevated in arteries in old age^5^. The accumulation of MGO-derived AGEs has been implicated in various pathophysiological processes, including excessive oxidative stress and cellular senescence, both of which contribute to age-related aortic stiffening^4,5,6^.

Molecularly, MGO preferentially reacts with arginine (Arg) residues to form specific advanced glycation end-products (AGEs), such as MGO-derived hydroimidazolone isomer (MGH-1), an established marker of MGO-induced glycation stress^7^. Elevated circulating concentrations of MGH-1 are strongly related to CVD risk and CV-related mortality in individuals with type 2 diabetes^8^.

In the present study, we used a series of complementary *in vivo, ex vivo,* and *in vitro* experimental approaches to determine the causal role of MGO-induced glycation stress in aortic stiffening and the putative underlying mechanisms mediating this response, including excessive oxidative stress and cellular senescence. Additionally, we explored the therapeutic potential of Gly-Low, a cocktail consisting of the natural compounds nicotinamide, pyridoxine, thiamine, piperine, and alpha-lipoic acid^9^, in mitigating aortic stiffening, oxidative stress and cellular senescence mediated by MGO-induced glycation stress.

## Materials and methods

### Animals

Young male C57BL/6 mice were acquired from Jackson laboratory and aged male C57BL/6 were acquired from the colony of the National Institute of Aging (maintained by Charles River, Wilmington, MA). Before starting the study, mice were given 4 weeks to get acclimated. During the entirety of the study mice were group-housed with a 12-h:12-h light:dark cycle and given ad libitum access to standard rodent chow (Teklad 7917; Envigo, Indianapolis, IN, USA). For the intervention, the mice received 1% MGO in drinking water. For the Gly-Low intervention mice were fed either a standard low-fat chow diet (21% fat (kcal), 60% carbohydrate (kcal) Envigo: TD.200743), a standard low-fat chow diet supplemented with our Gly-Low compound cocktail (21% fat (kcal), 60% carbohydrate (kcal) Envigo: TD.200742). Additional details regarding the Gly-Low cocktail, quantification of MGO and MGH-1 *in vivo* and *ex vivo*, other procedures, and antibodies/primer sequences used are provided in the supplemental methods.

### Aortic PWV

Aortic stiffness was assessed using the reference standard non-invasive *in vivo* measure, aortic pulse wave velocity (PWV), one week after the intervention, as previously described^12,31^. Briefly, mice were placed under light isoflurane anesthesia (1.0%–2.5%) and positioned supine on a warmed heat pad. Front- and hind-limb paws were then secured to corresponding ECG electrodes. Two Doppler probes were then placed on the skin at the transverse aortic arch and the abdominal aorta. Once clear R-waves were registered, three repeated 2-second ultrasound tracings were recorded and average pre-ejection time (i.e., time between the R-wave of the ECG to the foot of the Doppler signal) was determined for each location. To calculate aortic PWV, the distance between the two probes was divided by the difference between the transverse aortic arch and abdominal aorta pre-ejection times (time_abdominal_ – time_arch_) and is reported as centimeters/second (cm/s).

### Ex vivo aortic intrinsic mechanical wall stiffness (elastic modulus)

Two 1mm segments of thoracic aorta were collected at sacrifice from intervention-naïve male and female C57BL/6 mice and cleared of any perivascular connective tissue. Then, aortic ring from each donor animal were incubated in the following conditions in duplicate for 48 hours in various conditions. The control condition was DMEM + 1% penicillin-streptomycin (Standard media). For other agents used, their concentrations were 1uM TEMPOL, 500uM MGO, 100uM Gly-Low, 200nM MGH-1, 100uM Glo-1 inhibitor, and 1uM Doxo. After the incubation period, elastic modulus was assessed using the pin myograph (Danish Myo Technology, Denmark) exactly as described in the supplemental figure. Changes in aortic elastic modulus are reported as fold change relative to the control condition.

### Aortic RNA sequencing

RNA was isolated using Zymo research quick RNA miniprep kit (cat # 11-328) according to the manufacturer’s recommendations. Isolated RNA was sent for library preparation and sequencing by Novogene Corporation Inc. where RNA was poly-A selected using poly-T oligo-attached magnetic beads, fragmented, reverse transcribed using random hexamer primers followed by second strand cDNA synthesis using dTTP for non-directional library preparation. Samples underwent end repair, A-tailing, adapter ligated, size selected, amplified, and purified which is described in more detail in the supplemental methods.

### Statistical Analysis

Details of all the statistical analysis performed are provided in the supplemental methods. All data is presented as Means ± SEM in text and figures unless specified otherwise. Statistical significance was set to α=0.05. All statistical analyses were performed using Prism, version 10 (GraphPad Software, Inc, La Jolla, CA).

## Results

### Chronic MGO-induced glycation stress induces aortic stiffening in young mice

As a first step in elucidating the impact of MGO-induced glycation stress, we established a glycation stress model through the pharmacological administration of MGO in drinking water at a 1% concentration in young adult mice and compared them to young adult mice that received regular drinking water without MGO, as previously described^10^. This model allows for the simulation of chronic glycation stress in a controlled environment independent of aging. Following two months of daily MGO consumption, aortic PWV was measured *in vivo* to evaluate MGO-induced aortic stiffening (**Fig. 1a**). Aortic PWV is the gold-standard non-invasive approach for assessing aortic stiffness and is corollary to the reference standard measure of aortic stiffness in people, carotid-femoral PWV^11^. Chronic MGO consumption was associated with an ∼20% higher aortic PWV in young adult mice (MGO, 414 ± 21 vs. control, 342 ± 25 cm/sec; *P*=0.03) (**Fig 1b**). Additionally, MGO consumption caused a 2-fold elevation in circulating MGH-1 (*P*<0.001 vs. control) (**Fig. 1c**) and previous research has shown that Gly-Low supplementation significantly lowers circulating MGH-1 levels in young mice^9^. To determine if glycation stress-induced aortic stiffening can be prevented by inhibiting MGO, an additional group of mice were treated with glycation lowering compounds (Gly-Low) concurrently with MGO. We found that Gly-Low effectively prevented the MGO-induced increase in aortic PWV (MGO + Gly-low, 333 ± 6 cm/sec; *P* = 0.009 vs. MGO; *P*=0.93 vs. Control) (**Fig. 1b**).

**Figure 1:**
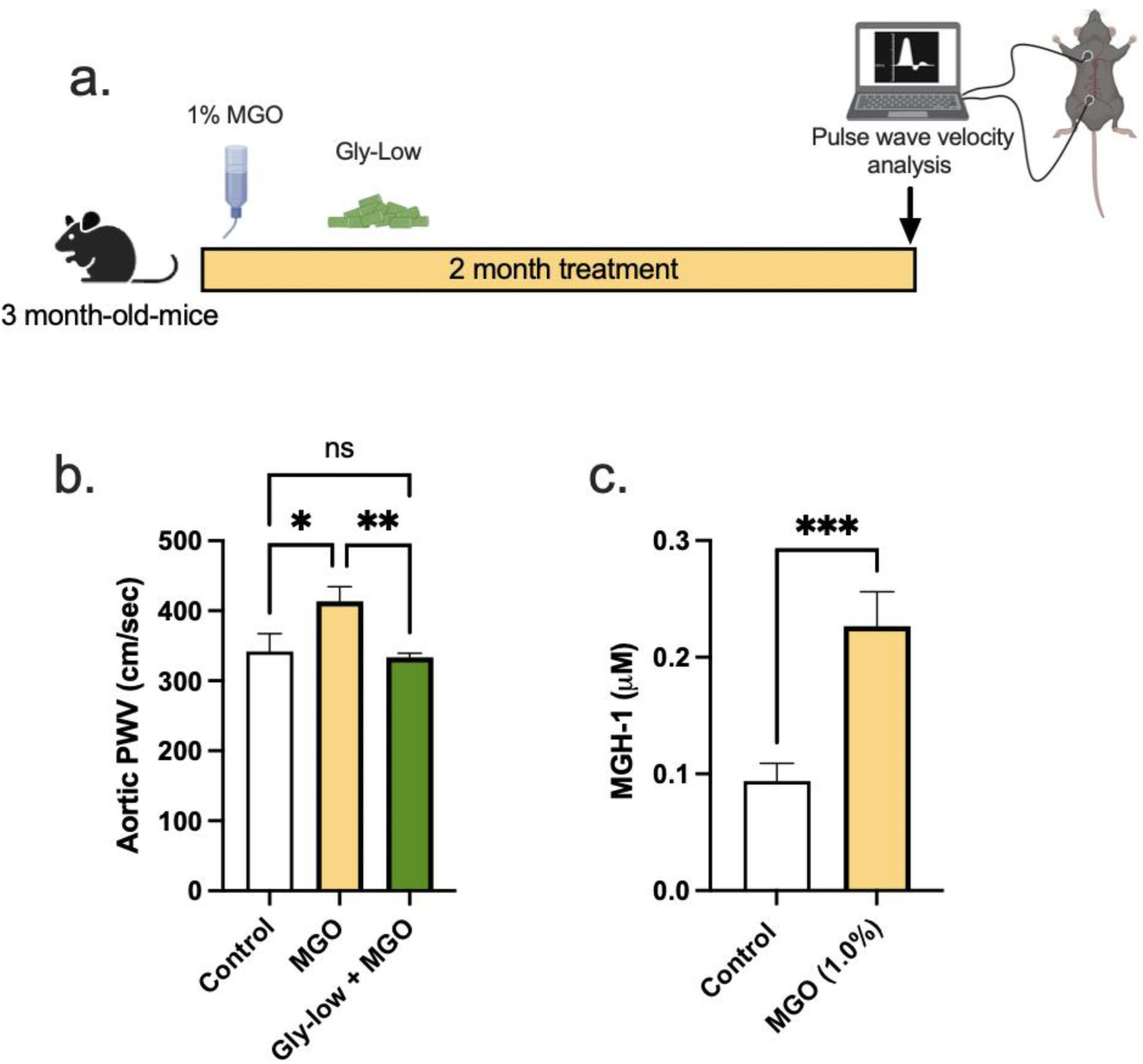
Methylglyoxal (MGO) induces glycation stress which can be prevented by Glycation lowering (Gly-Low) compound supplementation. (a) Treatment paradigm for MGO and Gly-low supplementation in 3-month-old mice. (b) Aortic PWV measured after intervention (n=7-10/group). (c) Plasma MGH-1 levels in mice that received normal chow with MGO supplemented in water (n=8-10). All values are mean ± SEM, *p<0.05

These findings emphasize the direct pathogenic role of MGO-induced glycation stress in aortic stiffening and illustrate the therapeutic potential of glycation-lowering compounds, like Gly-Low, in preventing aortic stiffening associated with glycation stress.

### Direct effect of MGO and Gly-Low on the ex-vivo stiffening on aorta rings

Next, to investigate the potential impact of MGO and Gly-Low on the intrinsic material stiffness of the aortic wall, we conducted *ex vivo* experiments using excised aorta rings from intervention-naive young adult (6 months old) male and female C57BL/6J mice and performed assessments of elastic modulus. Excised aorta rings from intervention-naïve young adult mice were incubated for 48 hours under the following conditions: (*1*) DMEM + 1% penicillin-streptomycin (control standard media), (*2*) Standard media + 500 μM MGO, (*3*) Standard media + 100 μM Gly-Low, and (*4*) Standard media + 500 μM MGO + 100 μM Gly-Low. Aortic elastic modulus assessments were subsequently performed as described previously^12^ (**Fig. 2a**). Aortic rings treated with MGO exhibited a 2-fold increase in elastic modulus (*P*=0.003 vs. control condition), indicative of increased intrinsic aortic wall stiffness (**Fig. 2b**). This MGO-induced aortic wall stiffening was attenuated by co-treatment with Gly-Low (*P=*0.009 MGO vs. MGO + Gly-Low) (**Fig. 2b**). Notably, Gly-Low treatment alone had no off-target effects on aortic elastic modulus (i.e., no difference between control and Gly-Low alone, *P*=0.40), confirming its specificity to MGO-induced glycation stress (**Fig. 2b**). MGO exposure increased the accumulation of the MGO-derived AGE, MGH-1 (*P*=0.004, MGO vs Control) (**Fig. 2c**), and co-exposure of MGO with Gly-Low markedly attenuated this increase in MGH-1 expression (P=0.04, MGO vs MGO+Gly-Low). The latter demonstrates molecular target engagement and further supporting its protective role against MGO-induced glycation stress (**Fig. 2c**). Together, these findings demonstrate that MGO-induced glycation stress directly increases aortic stiffness and promotes AGE accumulation in arteries, independent of other contributing *in vivo* factors.

**Figure 2:**
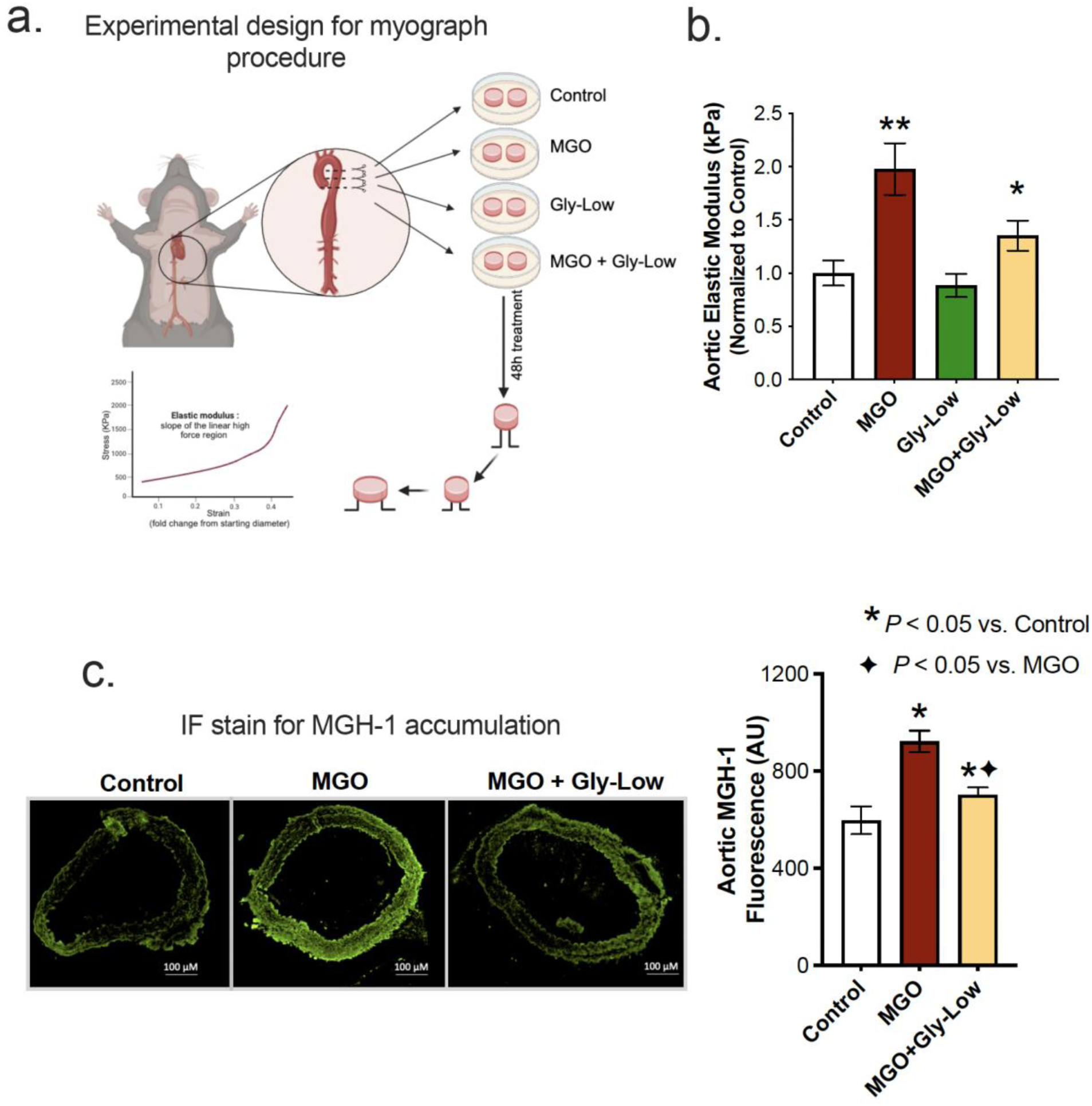
Ex-vivo incubation with MGO increases arterial stiffness which is mitigated by Gly-Low coincubation as measured by aortic elastic modulus. (a) Experimental design for the myograph procedure. (b) Aortic elastic modulus following exposure to standard media (control), MGO, Gly-Low, and MGO+Gly-Low (n=13/group). (c,d) Representative immunofluorescence staining for methylglyoxal derived hydroimidazolone-1 (MGH-1) accumulation in aortic rings exposed to standard media (control), MGO, and MGO+Gly-Low (n=5-8/group). All values are mean ± SEM, *p<0.05 vs control; ◆p<0.05 vs MGO alone.

### MGO-induced oxidative stress drives aortic stiffening

Elevated glycation stress has been shown to induce excessive reactive oxygen species (ROS)-related oxidative stress bioactivity in various cell types^13,14^. Considering the established role of excessive ROS bioactivity in aortic stiffening^6,15^, we next sought to investigate whether MGO-induced aortic stiffening is mediated through ROS-dependent mechanisms. To accomplish this, we first assessed superoxide bioactivity in aorta rings from intervention-naïve male and female mice via electron paramagnetic resonance spectroscopy (reference standard experimental approach for assessing ROS in biological tissues)^16^. Excised aorta rings were incubated for 48 hours under the following conditions: (*1*) Standard media (Control), (*2*) Standard media + 500 μM MGO, and (*3*) Standard media + 500 μM MGO + 100 μM Gly-Low. Incubation with MGO alone led to an ∼40% increase in aortic superoxide levels (*P*=0.04 vs. Control), and concomitant incubation of Gly-Low with MGO prevented this increase in superoxide bioactivity (*P*=0.03 MGO vs. MGO+GlyLow) (**Fig. 3a**).

**Figure 3:**
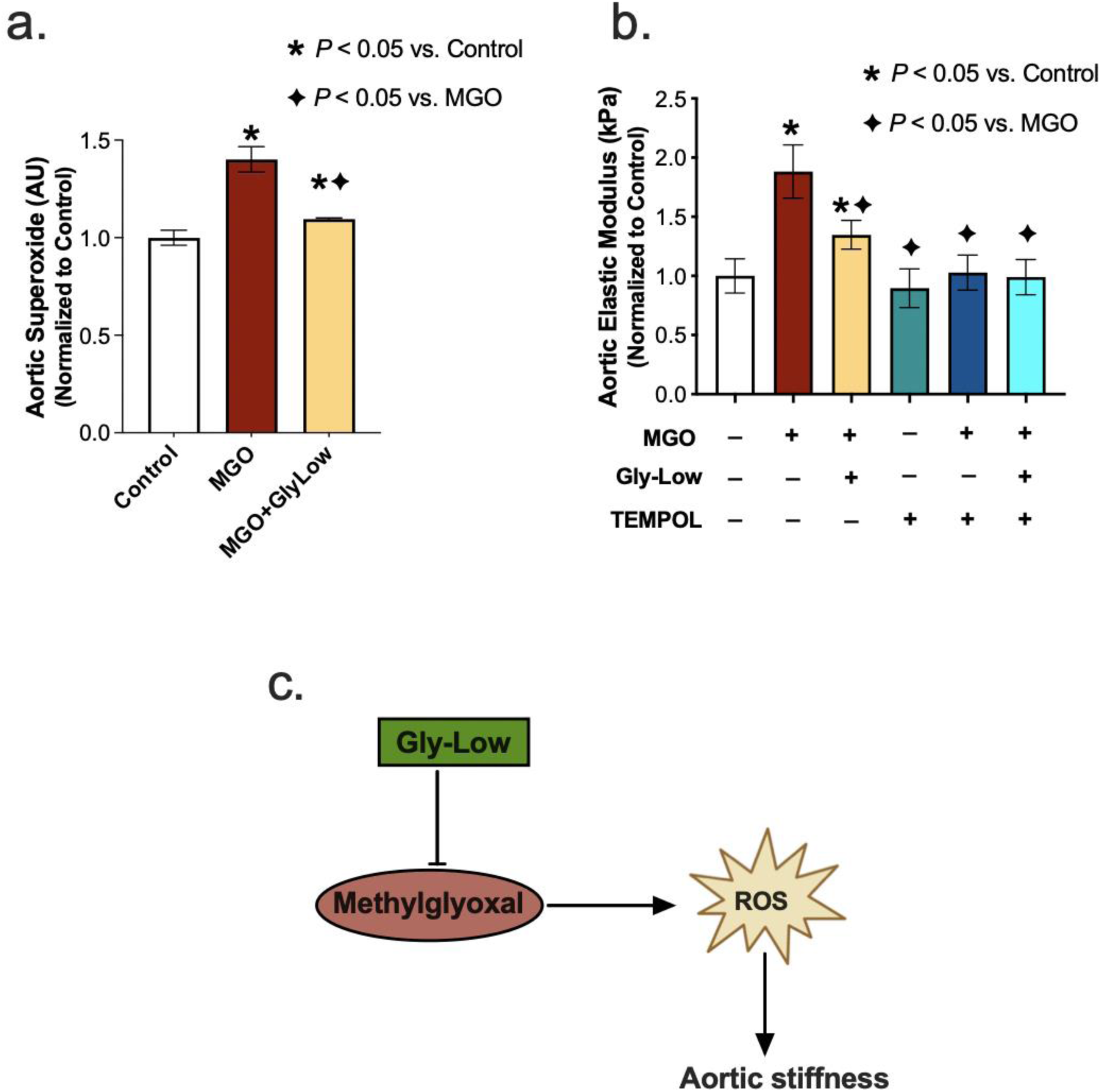
MGO-induced glycation stress drives aortic stiffness, in part, by increasing reactive oxygen species (ROS) levels. (a) Aortic superoxide levels assessed via Electron Paramagnetic Resonance-spectroscopy in aortic rings after 48-hour incubation with standard media (control), MGO, and MGO+Gly-Low (n=5/group) (b) Elastic modulus in aortic rings incubated with/without superoxide scavenger TEMPOL (n=8/group). (c) Mechanistic schematic of depicting MGO-induced glycation stress causes aortic stiffness via increasing ROS levels. All values are mean ± SEM, *p<0.05 vs control; ◆ p<0.05 vs MGO alone.

Next, to interrogate the direct role of excessive ROS bioactivity in aortic stiffening induced by MGO-mediated glycation stress, we incubated aorta rings from young adult intervention naïve mice for 48 hours with the superoxide scavenger, TEMPOL (1 μM), in the presence of MGO and Gly-Low, and assessed elastic modulus as described above. As previously demonstrated, incubation with MGO alone caused a 75% increase in elastic modulus compared to control condition (*P*=0.002), indicating MGO-induced aortic stiffening, which was fully prevented by coincubation with TEMPOL (*P*=0.01). Additionally, when aortic rings were incubated with MGO and Gly-Low, the increase in aortic stiffening was attenuated (corroborating our findings presented in Fig 2) (**Fig. 3b**). Moreover, the combined incubation with MGO, TEMPOL, and Gly-Low completely prevented the increase in stiffness, suggesting Gly-Low likely prevented MGO-mediated aortic stiffening by mitigating excessive ROS bioactivity (**Fig. 3b**). Notably, TEMPOL alone did not produce any off-target effects on elastic modulus, as evidenced by the minimal effects on the control conditions (**Fig. 3b**). These data suggest that MGO-induced glycation stress contributes to aortic stiffening, in part, through excessive ROS bioactivity (**Fig. 3c**), underscoring the potential of targeting glycation stress as a therapeutic strategy for mitigating excessive ROS-mediated aortic stiffening.

### MGH-1 Directly Induces Aortic Stiffening

MGH-1 is a prevalent AGE associated with various age-related disorders^7,8^. MGH-1, a non-crosslinking AGE, is predominantly produced from MGO and has been implicated in CVD^8,29,30^, but its direct effect on aortic stiffening is unknown. Thus, we aimed to determine if MGH-1 directly influences intrinsic mechanical wall stiffness of the aorta (elastic modulus) and whether Gly-Low could prevent this effect. To investigate this, aorta rings were excised from young intervention naïve mice for 48 hours under the following conditions: (*1*) Standard media (control), (*2*) Standard media + 200 nM MGH-1, and (*3*) Standard media+200 nM MGH-1+100 μM Gly-Low. Aortic elastic modulus assessments were subsequently performed as described above. Additionally, RNA sequencing was conducted on aortic segments incubated with MGH-1 and MGH-1 + Gly-Low to determine the molecular events in arteries altered by glycation stress and how these processes are influenced by a glycation stress lowering compound (**Fig. 4a**).

**Figure 4:**
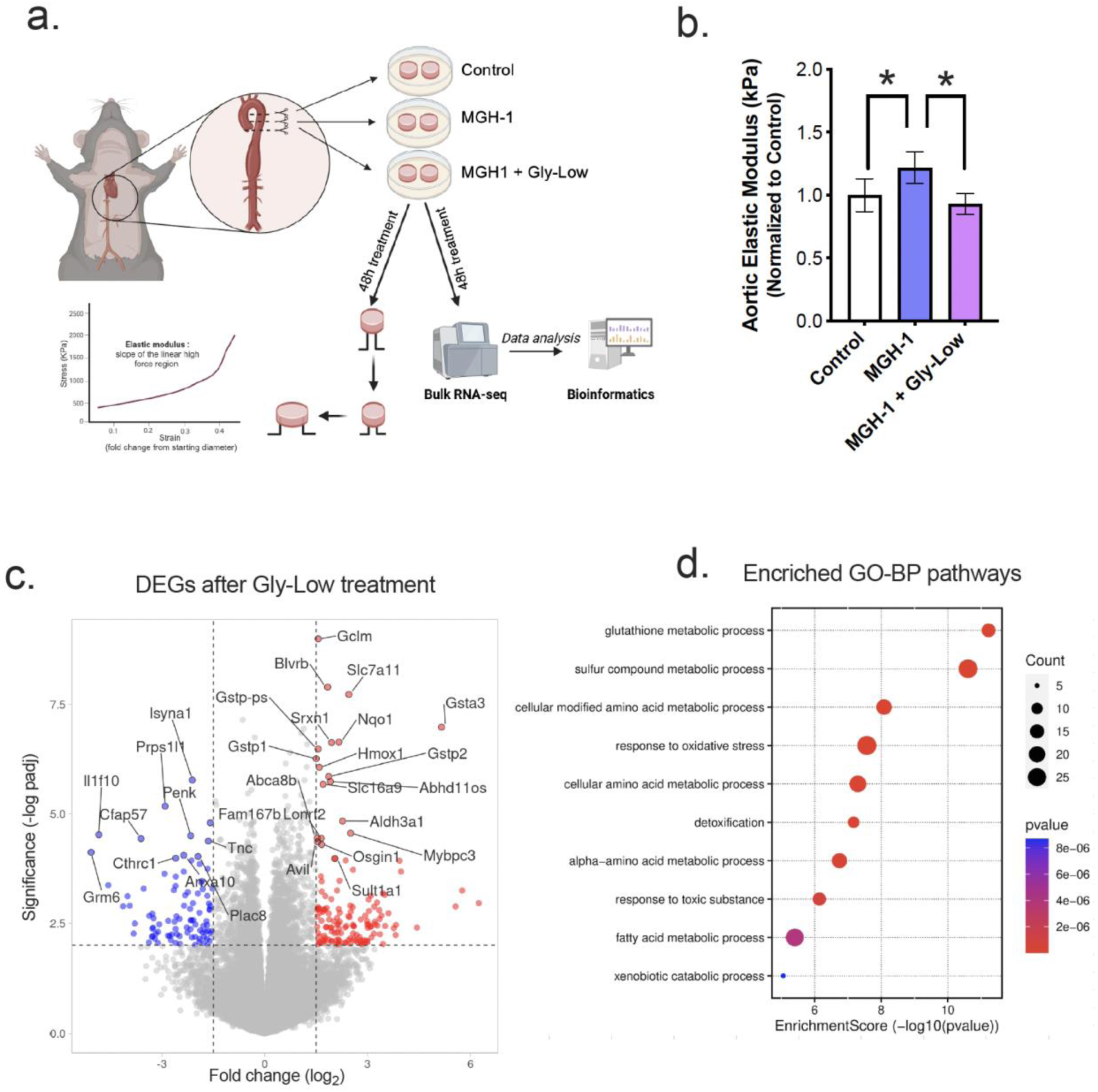
MGO specific isomer; MGH-1 can increase aortic stiffness in young mouse aortas and Gly-Low supplementation prevents it. (**a**) Study paradigm for the ex-vivo aortic stiffness measurement after 48-hour incubation with standard media (control), MGH-1, and MGH-1+Gly-Low. (**b**) Aortic elastic modulus for young (6-month) mouse aortic rings after incubation (n=9/group). (**c,d**) Differential gene expression and Gene ontology pathways in aortic rings incubated with MGH-1 and MGH-1+Gly-Low. All values are mean ± SEM, *p<0.05 vs control.

Following the 48-hour incubation period, incubation with MGH-1 increased (*P*=0.04 vs. control) aortic elastic modulus, an effect that was blocked by Gly-Low coincubation (*P*=0.02, MGH-1 vs. MGH-1+GlyLow) (**Fig. 4b**). Gene ontology pathway enrichment analysis of the differentially upregulated genes that were identified following bulk RNA sequencing indicated the activation of detoxification pathways and antioxidant responses with Gly-Low (**Fig. 4c-d**). The altered pathways suggest that Gly-Low may prevent aortic stiffening by mitigating excessive oxidative stress and these findings further connect its effects to the reduction in ROS bioactivity observed in our upstream *ex vivo* analyses.

### Methylglyoxal Induces Cellular Senescence in Endothelial Cells Independent of ROS Production

Endothelial cell senescence directly promotes aortic stiffening and vascular oxidative stress^12,22^. Given that MGO-mediated glycation stress induced aortic stiffening by increasing ROS bioactivity in the present study, and that cellular senescence has been implicated as a cause of excessive ROS production in endothelial cells and intact arteries^23–25^, we next hypothesized that glycation stress may induce cellular senescence in endothelial cells. To test this hypothesis, we treated HAECs with MGO and Doxo, which served as a positive control for senescence induction (**Fig. 5a**). We observed that MGO-derived glycation stress caused an increase in senescence biomarkers similar to Doxo, as evidenced by the increased intensity of senescence-associated β-galactosidase (SA-β-gal) (**Fig. 5b**) and higher expression of canonical senescence genes such as *Cdkn1a* (p21), *Cdkn2a* (p16) and *Serpine1* (PAI-1), and lower *Lmnb1* (Lambin B1) (**Fig. 5c**). In all conditions, lowering glycation stress via Gly-Low exposure generally mitigated the induction of cellular senescence (**Fig. 5b-c**). These findings indicated that MGO-induced glycation stress was a driver of cellular senescence in endothelial cells.

**Figure 5:**
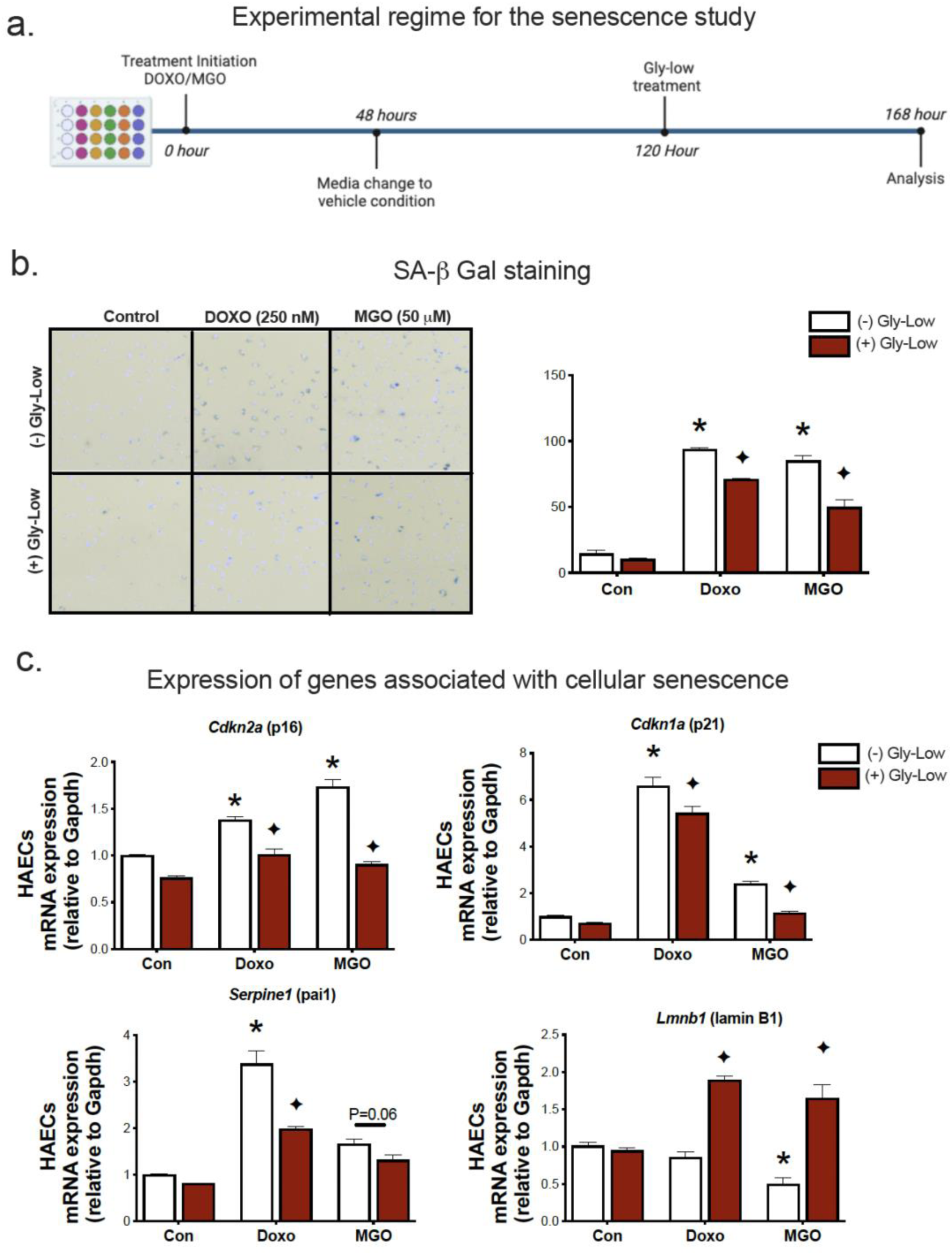
MGO-induced glycation stress can increase cellular senescence. (**a**) Study paradigm for the cell line treatment with DOXO/MGO and Gly-Low. (**b**) Senescence associated beta galactosidase staining for HAECs treated with MGO/DOXO with or without Gly-Low supplementation. (**c**) mRNA expression of various senescence associated genes. All values are mean ± SEM, *p<0.05 vs control; ◆ p<0.05 vs DOXO alone.

To further elucidate whether these senescence-promoting effects were mediated through ROS production, we treated HAECs with MGO in the presence and absence of the ROS scavenger TEMPOL. Our results demonstrated that TEMPOL treatment had no effect on MGO-induced cellular senescence in HAECs (**Fig. S1a-c**), implying that MGO induces cellular senescence in endothelial cells likely occurs through mechanisms that are independent of excessive ROS bioactivity and that cellular senescence likely precedes ROS production as a result of MGO-induced glycation stress.

### Gly-Low Mitigates Senescence-Induced Aortic Stiffening by Activating Glyoxalase-1 Mediated Detoxification Pathway

Preliminary studies from our laboratory have demonstrated that Doxo promotes aortic stiffening as a result of excessive cellular senescence^31^. Moreover, our RNA sequencing analysis indicated that Gly-Low may exert its anti-glycation stress properties by activating detoxification pathways. Thus, we next hypothesized that Gly-Low may mitigate the effects of senescence-induced (via Doxo exposure) aortic stiffening by activating the detoxification system. To test this hypothesis, we incubated excised aorta rings from young intervention naïve male and female mice for 48 hours under the following conditions: (*1*) Standard media (control), (*2*) Standard media+1µM Doxo, (*3*) Standard media+1 µM Doxo+100 μM Gly-Low, and (4) Standard media+100 μM Gly-Low. Doxo-incubated aorta rings exhibited higher aortic stiffness compared to the control condition (*P*=0.006). However, concomitant incubation of Doxo and Gly-Low prevented the increase in aortic stiffening observed with Doxo (*P*=0.02; Doxo + Gly-Low vs. Doxo alone) (**Fig 6a**), indicating that cellular senescence is likely a mechanism underlying glycation stress-induced aortic stiffening.

**Figure 6:**
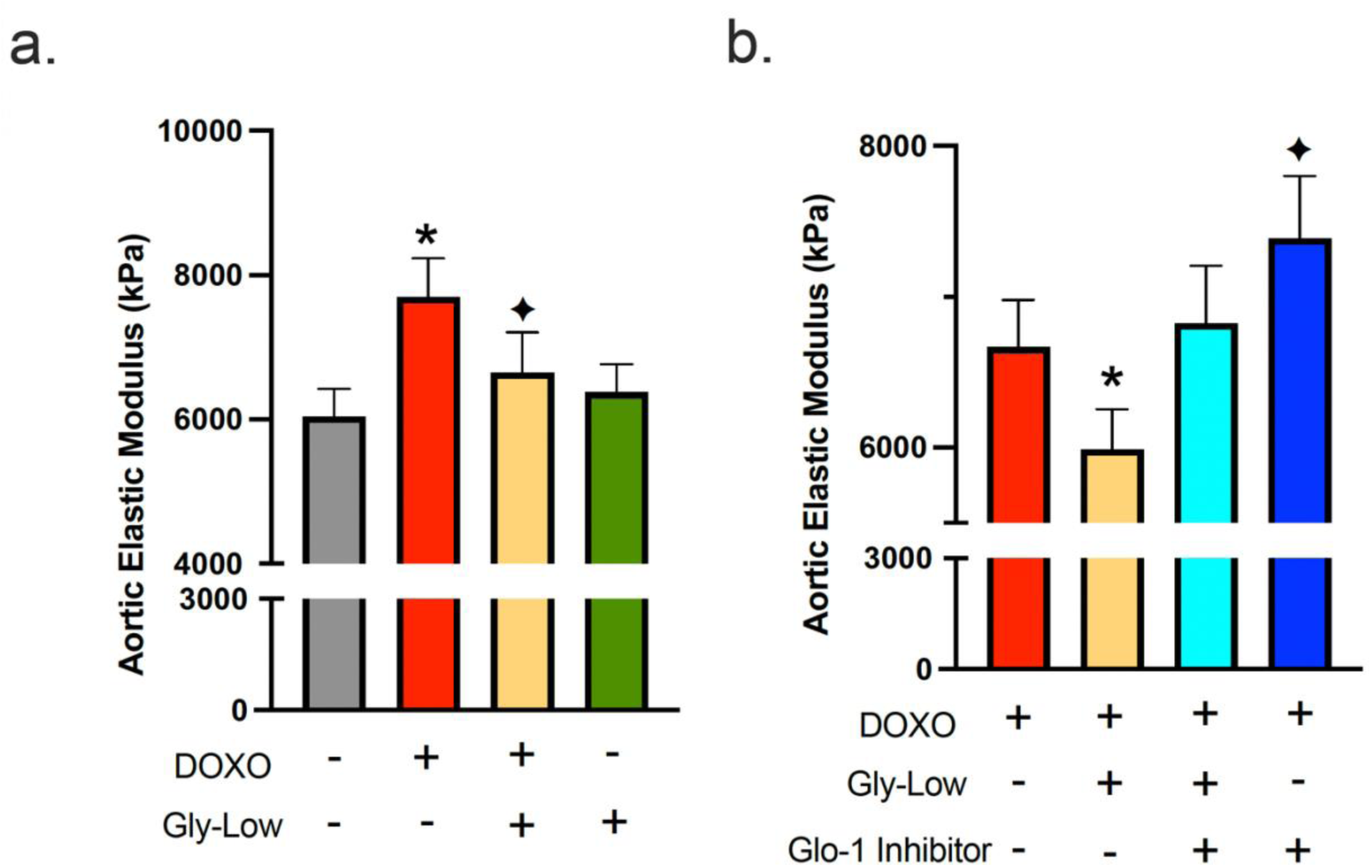
Gly-Low supplementation mitigates DOXO-induced aortic stiffening. (**a**) Aortic elastic modulus in young mouse aortas after 48-hour incubation with DOXO/Gly-Low (n=8/group). (**b**) Aortic elastic modulus in young mouse aortas after 48-hour incubation with DOXO/Gly-Low in absence/presence of Glo-1 inhibitor (n=9/group). All values are mean ± SEM, *p<0.05 vs control (No DOXO/No Gly-Low); ◆p<0.05 vs DOXO alone.

Considering the canonical role of the Glyoxalase-1 (Glo-1) detoxification system in clearing excessive MGO-induced glycation stress^10,18,19,20^, we next investigated whether the effects of Gly-Low were mediated through the Glo-1 signaling pathway. Aortic rings incubated with an established Glo-1 inhibitor^21^ in the presence of Doxo and Gly-Low showed no change in stiffness compared to the Doxo group alone (*P*=0.15). Additionally, aorta rings incubated with Doxo in the presence of the Glo-1 inhibitor exhibited significantly increased (*P*=0.03) aortic stiffness compared to Doxo alone (**Fig. 6b**). Collectively, these results suggest that Doxo-mediated aortic stiffening is induced by glycation stress, in part via suppression of Glo-1, and Gly-low mechanistically exerts its effect through activation of the Glo-1 pathway. Furthermore, our RNA sequencing data validates that Gly-Low mediates its therapeutic effect in part by activating the Glo-1 detoxification pathway. This suggests that activating or preserving Glo-1 may be a potential strategy for mitigating excessive cellular senescence-induced aortic stiffening.

### Reducing glycation stress may improve age-associated aortic stiffening

Building on our findings that MGO-induced glycation stress causally drives aortic stiffening independent of advanced age and given that glycation stress naturally accumulates with aging due to impaired activation of detoxification pathways (e.g., Glo-1)^5^, we next sought to determine whether reducing glycation stress through Gly-Low supplementation lowers aortic stiffness in old mice. To test this, we supplemented 20-month-old mice with Gly-Low in their drinking water (**Fig 7a**). After a four-month treatment period, we observed that Gly-Low-supplemented mice had lower aortic PWV than age-matched controls (Old Gly-Low, 352 ± 11 vs. Old control, 407 ± 12 cm/sec; *P*=0.01) (**Fig 7b**) and similar levels to that of young control mice (342 ± 25 cm/sec **Fig 1B**). Gly-Low supplementation also lowered levels of circulating MGO (*P*=0.002) and the MGO-derived AGE; MGH-1 (*P*=0.01), by ∼50% compared to age-matched control mice (**Fig. 7c-d**).

**Figure 7:**
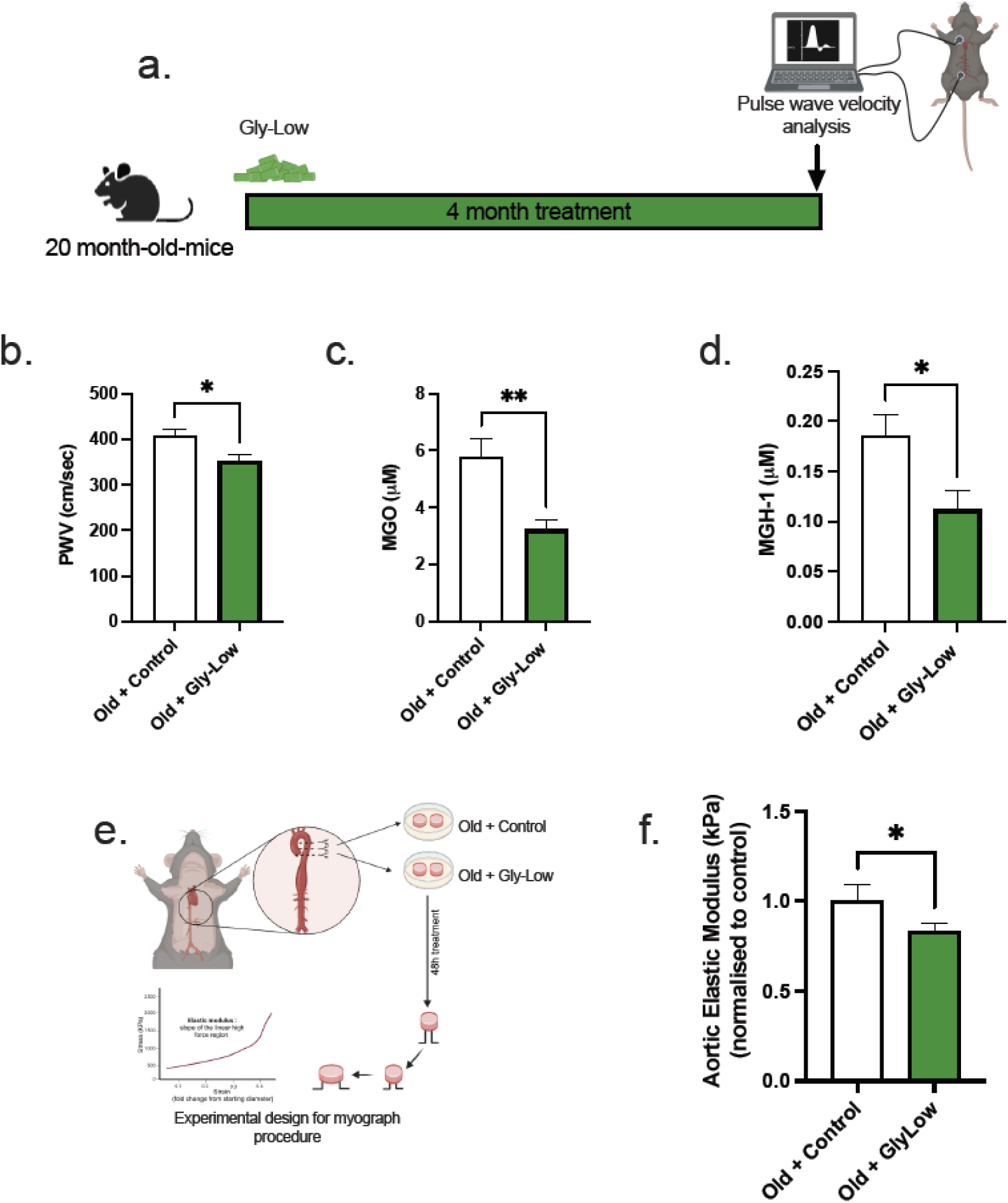
Gly-Low lowers MGO-induced glycation stress and aortic stiffening in old mice. (a) Study paradigm for the old mouse study. (b) Aortic stiffness as measured by PWV between two study groups (n=8-10). (c) MGO levels in plasma that received control and Gly-Low enriched diet (n=8-10). (d) Plasma MGH-1 levels (n=8-10). (e,f) Ex-vivo study paradigm focusing on aortic stiffness in old mouse aortas after 48-hour incubation in standard media (control) and Gly-Low (n=8). All values are mean ± SEM, *p<0.05 vs control

To further investigate the direct effect of Gly-Low on intrinsic mechanical wall stiffness (elastic modulus) in the setting of advanced age, we incubated aortic rings obtained from 24–26-month-old mice for 48 hours in the absence and presence of Gly-Low. Following the incubation, we assessed elastic modulus as described above (**Fig. 7e**). Our results indicated a significant reduction in stiffness in old mouse aorta rings incubated with Gly-Low (*P*=0.04), suggesting a direct role of Gly-Low in lowering aortic stiffness with aging (**Fig. 7f**). Collectively, these findings suggest that excessive glycation stress associated with aging directly contributes to age-related aortic stiffening.

## Discussion

In this study, we investigated the impact of MGO-induced glycation stress on aortic stiffness in young and old mice, explored the possible molecular mechanisms involved, and evaluated the therapeutic potential of the glycation-lowering compound, Gly-Low. Our findings provide novel and significant insights into the putative role of MGO-induced glycation stress as a mediator of age-related aortic stiffening and provide proof-of-principle efficacy for the use of glycation stress lowering compounds for mitigating aortic stiffening. Our results demonstrate that chronic MGO-induced glycation stress significantly increases aortic stiffness in both young and old mice. The observed impairments in aortic stiffness following MGO treatment underscore the critical role of AGEs in promoting age-related aortic stiffening. This effect was particularly pronounced in our pharmacological model of glycation stress, where young adult mice exhibited a marked increase in aortic stiffness, after just two months of MGO exposure. Interestingly, our study also reveals the direct impact of MGO on aortic stiffening, further supporting the notion that MGO-induced glycation stress can independently drive aortic stiffening. Moreover, our *ex vivo* experiments demonstrate that MGO directly increased aortic stiffness, accompanied by elevated AGE accumulation. These findings suggest that MGO-derived glycation stress directly drives AGE accumulation and functional alterations in the arterial wall. Moreover, our findings establish a putative role of excessive oxidative stress and cellular senescence in mediating aortic stiffening induced by MGO-derived glycation stress.

Key mechanistic insight from our study is the role of excessive ROS in mediating glycation stress-induced aortic stiffening. MGO treatment led to a substantial increase in aortic superoxide production, an effect that was significantly attenuated by Gly-Low supplementation. The use of the ROS scavenger TEMPOL further validated the involvement of oxidative stress in this process, as TEMPOL treatment effectively prevented MGO-induced aortic stiffening. This highlights the critical role of ROS in the pathogenesis of glycation stress-associated aortic stiffening. However, large-scale randomized controlled trials using chronic antioxidant supplementation have failed to demonstrate improvements in arterial health^26,27^, highlighting the need for alternative therapeutic strategies that mitigate excessive oxidative stress. Our findings suggest that targeting glycation stress may be a promising approach to mitigate excessive oxidative stress-related aortic stiffening. Moreover, our findings on the induction of cellular senescence by MGO provide additional insights into the molecular mechanisms by which glycation stress may promote aortic stiffening. MGO treatment of aortic endothelial cells induced a senescent phenotype characterized by increased SA-β-gal positive cells and elevated biomarkers of senescence. The absence of effect of TEMPOL on MGO-induced senescence indicates that senescence may be a driver of excessive ROS production in arteries (rather than vice versa) as a result of glycation stress. These finding highlight the intricate molecular-cellular processes underlying age-related aortic stiffening, suggesting an interactive relation between glycation stress, cellular senescence, ROS and aortic stiffening. The involvement of the Glo-1 detoxification pathway in the therapeutic effects of Gly-Low was another significant finding of our study. Our RNA sequencing data indicated that Gly-Low activates detoxification pathways, potentially contributing to the reduction of MGO-induced glycation stress. The inhibition of the Glo-1 pathway abrogated the beneficial effects of Gly-Low, confirming that this detoxification system is essential for mediating the protective actions of the compound against glycation-induced aortic stiffening.

The therapeutic potential of Gly-Low was evident in our *in vivo*, *ex vivo* and *in vitro* models. In aged mice, Gly-Low treatment significantly reduced MGO levels and MGO-derived AGEs, leading to a notable improvement in aortic stiffness. This suggests that Gly-Low effectively mitigates the adverse effects of glycation stress, likely by enhancing the detoxification of MGO and reducing the accumulation of pathological AGEs. The *ex vivo* experiments further confirmed the protective effects of Gly-Low, as co-incubation with Gly-Low prevented the MGO-induced increase in the elastic modulus of aortic rings. These findings underscore the potential of Gly-Low as a therapeutic agent for combating aortic stiffness associated with aging and glycation stress.

## CONCLUSION

Our study elucidates the putative molecular-cellular mechanisms by which MGO-induced glycation stress contributes to aortic stiffening and highlights the therapeutic potential of Gly-Low in mitigating these effects. By targeting glycation stress, reducing cellular senescence and oxidative stress, and activating detoxification pathways, Gly-Low offers a promising strategy for preserving arterial health and preventing the progression of CVD associated with aging or other conditions that may promote glycation stress in arteries (e.g., metabolic dysorders^5,28^). Further research is warranted to explore the clinical applicability of Gly-Low and other glycation-lowering compounds in managing arterial stiffness and improving CV outcomes in at-risk populations.

## Supporting information

Graphical abstract

## Author contributions

PS, RV, DRS, PK, and ZC, were involved in conceptualization and supervision. PS, RV, PK, and ZC drafted the manuscript. PS, RV, PK JG and ZC were involved in analysis and visualization. PS, RV, SAM, MAD, KRL, AZ, KK, LEN, LW, MT, MMS, EM, RS, ZC were involved in experimental procedures and live animal experimentation. All authors have read and agreed to the final version of the manuscript.

## Funding statement

We acknowledge support from the National Institute of Health: R01AG068288 (PK), R01AG061165 (PK), the Larry L. Hillblom Foundation (PS), NIH K99 HL159241 (ZC).

## Abbreviations

AGEs: Advanced glycation end products
Cdkn1a: Cyclin-dependent kinase inhibitor 1A (p21)
Cdkn2a: Cyclin-dependent kinase inhibitor 2A (p16)
CVDs: Cardiovascular diseases
DMEM: Dulbecco’s Modified Eagle Medium
Doxo: Doxorubicin
Glo-1: Glyoxalase-1
Gly-Low: Glycation-lowering compound
HAECs: Human aortic endothelial cells
Lmnb1: Lamin B1
MGH-1: Methylglyoxal-derived hydroimidazolone isomer-1
MGO: Methylglyoxal
PWV: Pulse wave velocity
ROS: Reactive oxygen species
SA-β-gal: Senescence-associated β-galactosidase
Serpine1: Plasminogen activator inhibitor-1 (PAI-1)
TEMPOL: 4-Hydroxy-TEMPO (a superoxide scavenger)

---

**Figure S1:**
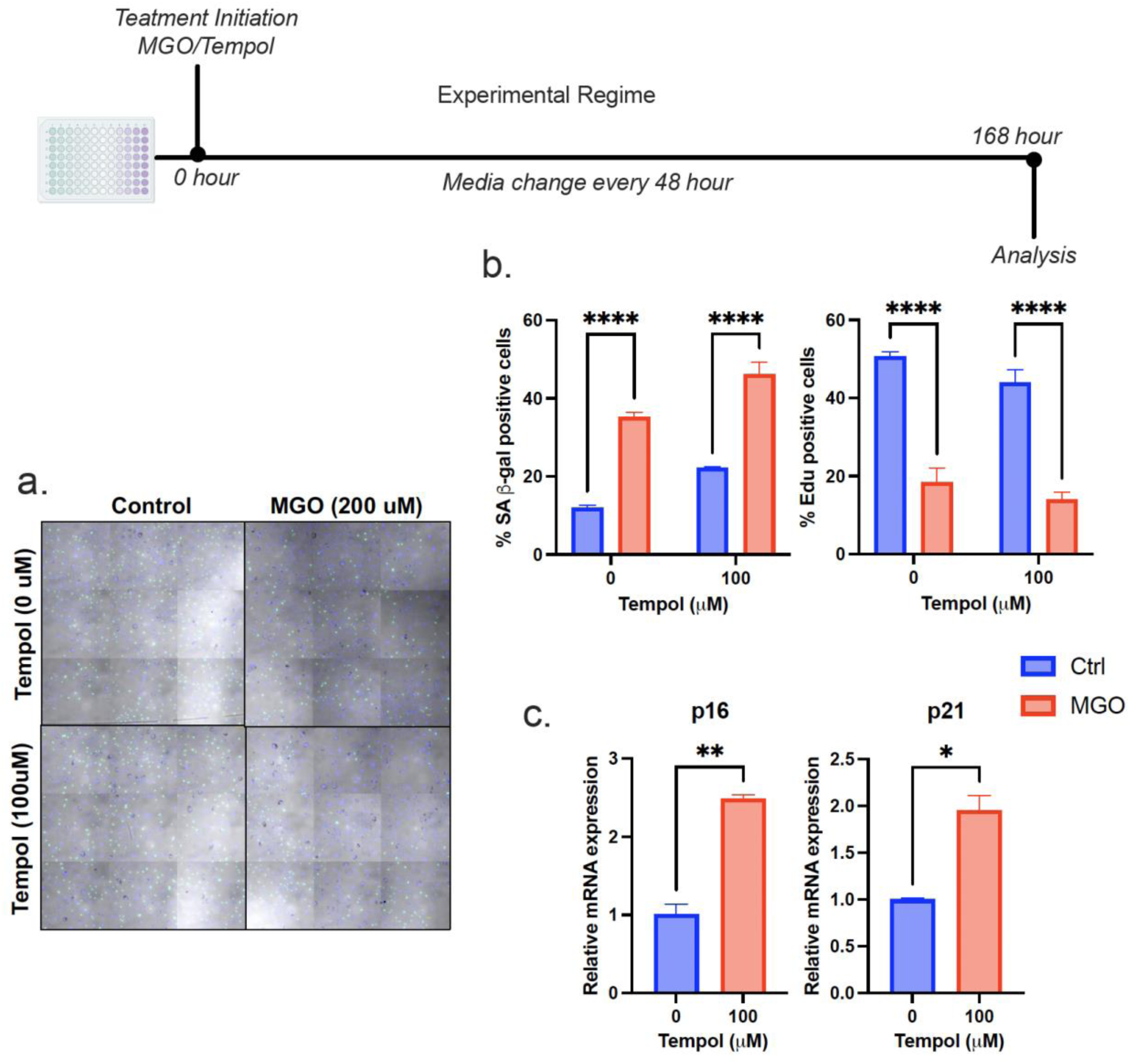
Methylglyoxal induces cellular senescence in endothelial cells independent of ROS levels. (**a**) Senescence associated beta galactosidase and EdU staining for HAECs treated with MGO/TEMPOL (**b**) Quantitative analysis of SA-beta and EdU staining. (c) mRNA expression of various senescence associated genes. All values are mean ± SEM, *p<0.05 vs control; ****p<0.0001 vs control.

## Notes

### Competing Interest Statement

The authors have declared no competing interest.

## References

1. Laurent, S. et al. Aortic stiffness is an independent predictor of all-cause and cardiovascular mortality in hypertensive patients. Hypertension 37, 1236–1241 (2001).

2. Lakatta, E. G. & Levy, D. Arterial and cardiac aging: major shareholders in cardiovascular disease enterprises: Part I: aging arteries: a “set up” for vascular disease. Circulation 107, 139–146 (2003).

3. Kopytek, M., Ząbczyk, M., Mazur, P., Undas, A. & Natorska, J. Accumulation of advanced glycation end products (AGEs) is associated with the severity of aortic stenosis in patients with concomitant type 2 diabetes. Cardiovasc. Diabetol. 19, 92 (2020).

4. Koyama, A. K., Pavkov, M. E., Wu, Y. & Siegel, K. R. Is dietary intake of advanced glycation end products associated with mortality among adults with diabetes? Nutr. Metab. Cardiovasc. Dis. 32, 1402–1409 (2022).

5. Chaudhuri, J. et al. The role of advanced glycation end products in aging and metabolic diseases: Bridging association and causality. Cell Metab. 28, 337–352 (2018).

6. Fleenor, B. S., Seals, D. R., Zigler, M. L. & Sindler, A. L. Superoxide-lowering therapy with TEMPOL reverses arterial dysfunction with aging in mice. Aging Cell 11, 269– 276 (2012).

7. McEwen, J. M., Fraser, S., Guir, A. L. S., Dave, J. & Scheck, R. A. Synergistic sequence contributions bias glycation outcomes. Nat. Commun. 12, 3316 (2021).

8. Hanssen, N. M. J. et al. Higher plasma methylglyoxal levels are associated with incident cardiovascular disease and mortality in individuals with type 2 diabetes. Diabetes Care 41, 1689–1695 (2018).

9. Wimer, L., et al. Combination therapy of glycation lowering compounds reduces caloric intake, improves insulin sensitivity, and extends lifespan. bioRxiv (2022) doi:10.1101/2022.08.10.503411.

10. Zunkel, K., Simm, A. & Bartling, B. Long-term intake of the reactive metabolite methylglyoxal is not toxic in mice. Food Chem. Toxicol. 141, 111333 (2020).

11. Chirinos, J. A., Segers, P., Hughes, T. & Townsend, R. Large-artery stiffness in health and disease: JACC state-of-the-art review. J. Am. Coll. Cardiol. 74, 1237– 1263 (2019).

12. Clayton, Z. S. et al. Cellular senescence contributes to large elastic artery stiffening and endothelial dysfunction with aging: Amelioration with senolytic treatment. Hypertension 80, 2072–2087 (2023).

13. Seo, K., Ki, S. H. & Shin, S. M. Methylglyoxal induces mitochondrial dysfunction and cell death in liver. Toxicol. Res. 30, 193–198 (2014).

14. Chan, C.-M. et al. Methylglyoxal induces cell death through endoplasmic reticulum stress-associated ROS production and mitochondrial dysfunction. J. Cell. Mol. Med. 20, 1749–1760 (2016).

15. Zhou, R.-H. et al. Mitochondrial oxidative stress in aortic stiffening with age: the role of smooth muscle cell function. Arterioscler. Thromb. Vasc. Biol. 32, 745–755 (2012).

16. Murphy, M. P. et al. Guidelines for measuring reactive oxygen species and oxidative damage in cells and in vivo. Nat. Metab. 4, 651–662 (2022).

17. Rabbani, N. et al. Glycation of LDL by methylglyoxal increases arterial atherogenicity: a possible contributor to increased risk of cardiovascular disease in diabetes. Diabetes 60, 1973–1980 (2011).

18. Peters, A. S. et al. Gender difference in glyoxalase 1 activity of atherosclerotic carotid artery lesions. J. Vasc. Surg. 62, 471–476 (2015).

19. Aragonès, G. et al. The glyoxalase system in age-related diseases: Nutritional intervention as anti-ageing strategy. Cells 10, 1852 (2021).

20. Chaudhuri, J. et al. A Caenorhabditis elegans model elucidates a conserved role for TRPA1-Nrf signaling in reactive α-dicarbonyl detoxification. Curr. Biol. 26, 3014– 3025 (2016).

21. McMurray, K. M. J. et al. Glo1 inhibitors for neuropsychiatric and anti-epileptic drug development. Biochem. Soc. Trans. 42, 461–467 (2014).

22. Jia, G., Aroor, A. R., Jia, C. & Sowers, J. R. Endothelial cell senescence in aging-related vascular dysfunction. Biochim. Biophys. Acta Mol. Basis Dis. 1865, 1802– 1809 (2019).

23. Mahoney, S. A. et al. Intermittent supplementation with fisetin improves arterial function in old mice by decreasing cellular senescence. Aging Cell 23, e14060 (2024).

24. Davalli, P., Mitic, T., Caporali, A., Lauriola, A. & D’Arca, D. ROS, cell senescence, and novel molecular mechanisms in aging and age-related diseases. Oxid. Med. Cell. Longev. 2016, 3565127 (2016).

25. Roger, L., Tomas, F. & Gire, V. Mechanisms and regulation of cellular senescence. Int. J. Mol. Sci. 22, 13173 (2021).

26. Biesalski, H. K., Grune, T., Tinz, J., Zöllner, I. & Blumberg, J. B. Reexamination of a meta-analysis of the effect of antioxidant supplementation on mortality and health in randomized trials. Nutrients 2, 929–949 (2010).

27. Sugamura, K. & Keaney, J. F., Jr. Reactive oxygen species in cardiovascular disease. Free Radic. Biol. Med. 51, 978–992 (2011).

28. Taniguchi, N. et al. Involvement of glycation and oxidative stress in diabetic macroangiopathy. Diabetes 45 Suppl 3, S81–3 (1996).

29. Kilhovd BK, Juutilainen A, Lehto S, et al. Increased serum levels of methylglyoxal-derived hydroimidazolone-AGE are associated with increased cardiovascular disease mortality in nondiabetic women. Atherosclerosis. 2009;205(2):590–594. doi:10.1016/j.atherosclerosis.2008.12.041

30. Ishibashi Y, Matsui T, Nakamura N, Sotokawauchi A, Higashimoto Y, Yamagishi SI. Methylglyoxal-derived hydroimidazolone-1 evokes inflammatory reactions in endothelial cells via an interaction with receptor for advanced glycation end products. Diab Vasc Dis Res. 2017;14(5):450–453. doi:10.1177/1479164117715855

31. Venkatasubramanian R, Mahoney SA, Maurer G, Darrah M, Ludwig K, VanDongen NS, Rossman MJ, Brunt VE, Campisi J, Seals DR and Clayton ZS, 2023. Senolytic administration following doxorubicin chemotherapy prevents large elastic artery stiffening and endothelial dysfunction. Physiology. 2023.doi: 10.1152/physiol.2023.38.S1.5733334

