## Supplementary material for "Methylglyoxal-induced glycation stress promotes aortic stiffening: Putative mechanistic roles of oxidative stress and cellular senescence": Methods

### Supplemental methods

#### Ethical approval

All protocols and procedures for this study were submitted to and approved by the University of Colorado Boulder (protocol #2618) and Buck Institute Research of Aging (protocol #A10249), Institutional Animal Care and Use Committee and adhered to the National Institutes of Health's Guide for the Care and Use of Laboratory Animals.

#### Study design and experimental animals

Young male C57BL/6 mice were acquired from Jackson laboratory and aged male C57BL/6 were acquired from the colony of the National Institute of Aging (maintained by Charles River, Wilmington, MA). Before starting the study, mice were given 4 weeks to get acclimated. During the entirety of the study mice were group-housed with a 12-h:12-h light:dark cycle and given ad libitum access to standard rodent chow *(*Teklad 7917; Envigo, Indianapolis, IN, USA*)*. For the intervention, the mice received 1% MGO in drinking water. For the Gly-Low intervention mice were fed either a standard low-fat chow diet (21% fat (kcal), 60% carbohydrate (kcal) Envigo: TD.200743), a standard low-fat chow diet supplemented with our Gly-Low compound cocktail (21% fat (kcal), 60% carbohydrate (kcal) Envigo: TD.200742). A combination of supplemental grade compounds, safe to be consumed in set dosages, were prepared and incorporated into a modified pre-irradiated standard AIN-93G mouse chow diet from Envigo. The cocktail consists of alpha lipoic acid (20.19%), nicotinamide (57.68%), thiamine hydrochloride (4.04%), piperine (1.73%), and pyridoxamine dihydrochloride (16.36%), and is supplemented in the diet to achieve a daily consumption rate in mg/kg of body weight/day. For diet, these percentages translate to 3 g/kg alpha lipoic acid, 8.57 g/kg nicotinamide, 0.6 g/kg thiamine hydrochloride, 0.26 g/kg piperine, and 2.43 g/kg pyridoxamine dihydrochloride.

#### *In Vivo Aortic Stiffness (PWV)*

Aortic stiffness was assessed using the reference standard non-invasive *in vivo* measure, aortic pulse wave velocity (PWV), one week after the intervention, as previously described[1,2].Briefly, mice were placed under light isoflurane anesthesia (1.0%–2.5%) and positioned supine on a warmed heat pad. Front- and hind-limb paws were then secured to corresponding ECG electrodes. Two Doppler probes were then placed on the skin at the transverse aortic arch and the abdominal aorta. Once clear R-waves were registered, three repeated 2-second ultrasound tracings were recorded and average pre-ejection time (i.e., time between the R-wave of the ECG to the foot of the Doppler signal) was determined for each location. To calculate aortic PWV, the distance between the two probes was divided by the difference between the transverse aortic arch and abdominal aorta pre-ejection times ($time_{abdominal} – time_{arch}$) and is reported as centimeters/second (cm/s).

***Sacrifice and tissue collection***.

Mice were sacrificed using a method approved under the American Veterinary Medical Association guidelines. Mice were anesthetized under inhaled anesthesia (open-drop method) and euthanized via cardiac exsanguination. The heart was removed, cleaned, and weighed. The aorta was excised and rinsed in physiological saline solution (PSS), cleared of perivascular adipose tissue, and sectioned and stored as described below.

***Ex vivo aortic intrinsic mechanical wall stiffness (elastic modulus)***.

Two 1mm segments of thoracic aorta were collected at sacrifice from intervention-naïve male and female C57BL/6 mice and cleared of any perivascular connective tissue. Then, aortic ring from each donor animal were incubated in the following conditions in duplicate for 48 hours in various conditions. The control condition was DMEM + 1% penicillin-streptomycin (Standard media). For other agents used, their concentrations were 1uM TEMPOL, 500uM MGO, 100uM Gly-Low, 200nM MGH-1, 100uM Glo-1 inhibitor, and 1uM Doxo. After the incubation period, elastic modulus was assessed using the pin myograph (Danish Myo Technology, Denmark) as previously described[1]. Briefly, aorta rings were mounted on to two prongs in a phosphate-buffered saline (PBS) bath heated up to 37C. After three rounds of pre-stretching (pins displaced to 1mm), the aortic diameter was increased until a force of 1mN was reached and then was incrementally increased by 50µM every three minutes until mechanical failure. The force corresponding to each stretching interval was recorded and utilized to calculate stress and strain and to construct a stress-strain curve:

$$\text{Strain } (\lambda) = \Delta d/d_i$$

Where d is diameter and $d_i$ is the initial diameter.

$$\text{Stress}(t) = (\lambda L)/2(HD)$$

Where L is one-dimensional load, H is intima-media thickness, and D is vessel length. The elastic modulus of the stress-strain curve was determined as the slope of the linear regression fit of the final four points of the stress-strain curve before the aortic rings broke as previously reported by our laboratory. Aortic wall thickness and diameter were assessed in 1mm aortic rings frozen in optimal cutting temperature, which were stored in -80C until the time of sectioning. Aortic sectioning was carried out in a cryostat (Leica biosystems, Weltzar, Germany) at -22°C. 7um sections were visualized and imaged under brightfield microscope and ImageJ was used to quantify the aortic thickness and diameter. Changes in aortic elastic modulus were presented as a fold-change relative to the control condition.

***Aortic ROS Levels***.

Whole-cell aortic ROS production was assessed by electron paramagnetic resonance (EPR) spectrometry using the spin probe 1-hydroxy-3-methoxycarbonyl-2,2,5,5-tetramethylpyrrolidine (CMH; Enzo Life Sciences, Farmingdale, NY, Cat. No. ALX-430-078), as previously described[1]. In short, two 1-mm aortic rings were washed in warm physiological saline solution and incubated in Krebs/HEPES buffer, consisting of 99 mM NaCl, 4.7 mM KCl, 1.87 mM CaCl2, 1.2 mM MgSO4, 25 mM NaHCO3, 1.03 mM KH2PO4, 20 mM Na-HEPES, 11.1 mM glucose, 0.1 mM diethylenetriaminepenta-acetic acid, 0.0035 mM sodium diethyldithiocarbamate, and Chelex (Sigma-Aldrich, Cat. No. C7901), containing 0.5 mM CMH at 37°C for 60min. Samples were analyzed using aMS300 Xband EPR spectrometer (Magnettech, Berlin, Germany) with the following instrument parameters: B0-Field, 3350G; sweep, 80G; sweep time, 60s; modulation, 3000mG; MWatten, 7dB; gain, 500.

***Aortic Immunofluorescence.***

Immunofluorescence assays were performed to visualize the subcellular localization of a prominent MGO byproduct methylglyoxal derived hydroimidazolone-1 (MGH-1). After the 48-hour incubation period mentioned in the subsection “Ex vivo aortic intrinsic mechanical wall” ~1mm section of thoracic aorta was excised and frozen in OCT (Tissue-Tek® O.C.T.) compound, as described previously[1]. Later in time, 7 µm sections (Leica CM1520) were plated on poly-L-lysine-coated microscope slides, fixed in 4% paraformaldehyde, washed with PBS, and permeabilized (0.1% Triton X-100). Slides were then stained with anti-MGH-1 primary antibody (1:200; Cell Biolabs, San Diego, CA; Cat# STA-011) according to instructions from the mouse-on-mouse immunodetection kit (Vector laboratories, Newark, CA; Cat# BMK-2202), and then incubated with a species-specific fluorescent secondary antibody (AlexaFluor 647; Invitrogen, Waltham, MA) for 30 minutes. Slides were washed, stained with DAPI (1:1000; Invitrogen, Waltham, MA; Cat# D1306) for 5 minutes, and cured overnight with ProLong Gold mounting media (Invitrogen, Waltham, MA; Cat# P36980). These slides were then imaged using EVOS m7000 (ThermoFisher, Waltham, MA; Cat# AMF7000) fluorescence microscope under identical conditions and analyzed using Invitrogen Celleste 5.0 Image Analysis Software. Images were obtained at 10x magnification. The abundance of MGH-1 was determined as the average intensity (A.U.) of the positively stained area across N = 6-8 samples/animal.

***HAEC RNA extraction and cDNA synthesis***

mRNA gene expression was measured in segments of thoracic aorta following mechanical homogenization. RNA was extracted using the RNeasy mini kit (Qiagen, Hilden, Germany). cDNA was synthesized using the iScript cDNA synthesis kit (Bio-Rad Laboratories, Hercules, CA). Transcripts of cellular senescence-related genes (primers reported in **Table S**) were analyzed using a StepOnePlus Real-Time PCR System (Applied Biosystems, Waltham, MA) in 96-well plates and the Taqman OpenArray (Applied Biosystems, Waltham, MA) was used as a master mix, as described[1,2] SimpleSeq DNA sequencing (Quintara Biosciences, Cambridge, MA) was used to validate PCR products.

Table of primer sequences for supplemental data.

| Gene | Species | Forward primer | Reverse primer |
|---|---|---|---|
| ***Cdkn2a*** | **Mouse** | **CCCAACGCCCCGAACT** | **GCAGAAGAGCTGCTACGTGAA** |
| ***Cdkn1a*** | **Mouse** | **TTGCCAGCAGAATAAAAGGTG** | **TTTGCTCCTGTGCGGAAC** |
| ***Serpine1*** | **Mouse** | **TGGAAGGGCAACATGACCAG** | **TCAGGCATGCCCAACTTCTC** |

| | | | |
|---|---|---|---|
| ***Lmnb1*** | **Mouse** | **GAGCCCCAAGAGCATCCAAT** | **CTGAGAAGGCTCTGCACTGT** |
| ***Gapdh*** | **Mouse** | **AAGGTCATCCCAGAGCTGAA** | **CTGCTTCACCACCTTCTTGA** |

***Aortic RNA sequencing***

RNA was isolated using Zymo research quick RNA miniprep kit (cat # 11-328) according to the manufacturer's recommendations. Isolated RNA was sent for library preparation and sequencing by Novogene Corporation Inc. where RNA was poly-A selected using poly-T oligo-attached magnetic beads, fragmented, reverse transcribed using random hexamer primers followed by second strand cDNA synthesis using dTTP for non-directional library preparation. Samples underwent end repair, A-tailing, adapter ligated, size selected, amplified, and purified. Illumina libraries were quantified using Qubit and qPCR and analyzed for size distribution using a bioanalyzer. Libraries were pooled and sequenced on an Illumina Novoseq 6000 to acquire paired-end 150 bp reads. Data quality was assessed, and adaptor reads, and low-quality reads were removed. Reads that passed the quality filtering process were mapped paired end to the reference genome (GRCm38) using Hisat2 v2.0.5. feature Counts v1.5.0-p3 was used to count reads that mapped to each gene. Differential expression analysis was performed using DESEq2 (1.20.0). Where indicated, bootstrapping was performed using R (R version 4.1.2) program 'boot' (1.3-28.1). To determine the expected mean and standard deviation, n=i log2 fold changes were randomly selected 1000 times, in which i is the number of genes in the gene set.

***Quantification of dicarbonyl***

200 µL of 80:20 MeOH:dd$H_2O$ (−80°C) containing 50 pmol $^{13}C_3$-MGO was added to 10 µL of serum and extracted at −80°C overnight. Insoluble protein was removed via centrifugation at 14,000 x g for 10 min at 4°C. Supernatants were derivatized with 10 µL of 10 mM o-phenylenediamine for 2 h with end-over-end rotation protected from light. Derivatized samples were centrifuged at 14,000 x *g* for 10 min, and the supernatant was chromatographed using a Shimadzu LC system equipped with a 150 x 2mm, 3µm particle diameter Luna $C_{18}$ column (Phenomenex, Torrance, CA) at a flow rate of 0.450 mL/min. Buffer A (0.1% formic acid in $H_2O$) was held at 90% for 0.25 min, then a linear gradient to 98% solvent B (0.1% formic acid in acetonitrile) was applied over 4 min. The column was held at 98% B for 1.5 min, washed at 90% A for 0.5 min, and equilibrated to 99% A for 2 min. Multiple reaction monitoring (MRM) was conducted in positive ion mode using an AB SCIEX 4500 QTRAP with the following transitions: *m/z*145.1→77.1 (MGO); *m/z* 235.0→157.0 (3-DG); m/z 131.0→77.0 (GO); m/z 161.0→77.0 (HPA); m/z 148.1→77.1 ($^{13}C_3$-MGO, internal standard).

***Quantification of free MGH-1***

10 µL of serum was added to 200 µL of 80:20 MeOH:dd$H_2O$ (−80°C) containing ten pmol $^{13}$C-MG-H1 and extracted at −80°C overnight. Insoluble protein was removed via centrifugation at 14,000 x g for 10 min at 4°C, and supernatants were transferred to a new tube. 15 µL of heptafluorobutyric acid (1:1 in $H_2O$) was added to each sample, and

debris was removed via centrifugation at 14,000 × *g* for 10 min. Samples were analyzed as described previously[3]. (QuARKMod).

#### *Statistical Analyses*

Statistical analyses were conducted using GraphPad Prism version 10 (GraphPad Software, Inc., San Diego, CA, USA; RRID:SCR_002798). Data were assessed for statistical outliers (ROUT test; Q = 1%), and outliers were excluded from final analyses. Differences in pulse wave velocity were assessed using two-way analysis of variance (ANOVA) with Šidák's post-hoc test when a main effect was observed. Differences across animal characteristics, reactive oxygen species production, were assessed using a paired T-Test. Differences in immunofluorescence intensity were assessed using paired *t*-test with the comparisons. All the data are presented as mean ± SEM. P values less than 0.05 were considered statistically significant. Significant differences are indicated: * $p \leq 0.05$, ** $p \leq 0.005$, *** $p \leq 0.0005$, **** $p < 0.0001$.

References


1. Clayton, Z. S. *et al.* Cellular senescence contributes to large elastic artery stiffening and endothelial dysfunction with aging: Amelioration with senolytic treatment. *Hypertension* **80**, 2072–2087 (2023)
2. Mahoney, S. A. *et al.* Intermittent supplementation with fisetin improves arterial function in old mice by decreasing cellular senescence. *Aging Cell* **23**, e14060 (2024).
3. Wimer .L. *et al*. Glycation-Lowering Compounds Inhibit Ghrelin Signaling to Reduce Food Intake, Lower Insulin Resistance, and Extend Lifespan. Bioxiv. 10.2139/ssrn.4471671 (2023).
