## Supplementary figures and images for "Methylglyoxal-induced glycation stress promotes aortic stiffening: Putative mechanistic roles of oxidative stress and cellular senescence"

### Graphical abstract

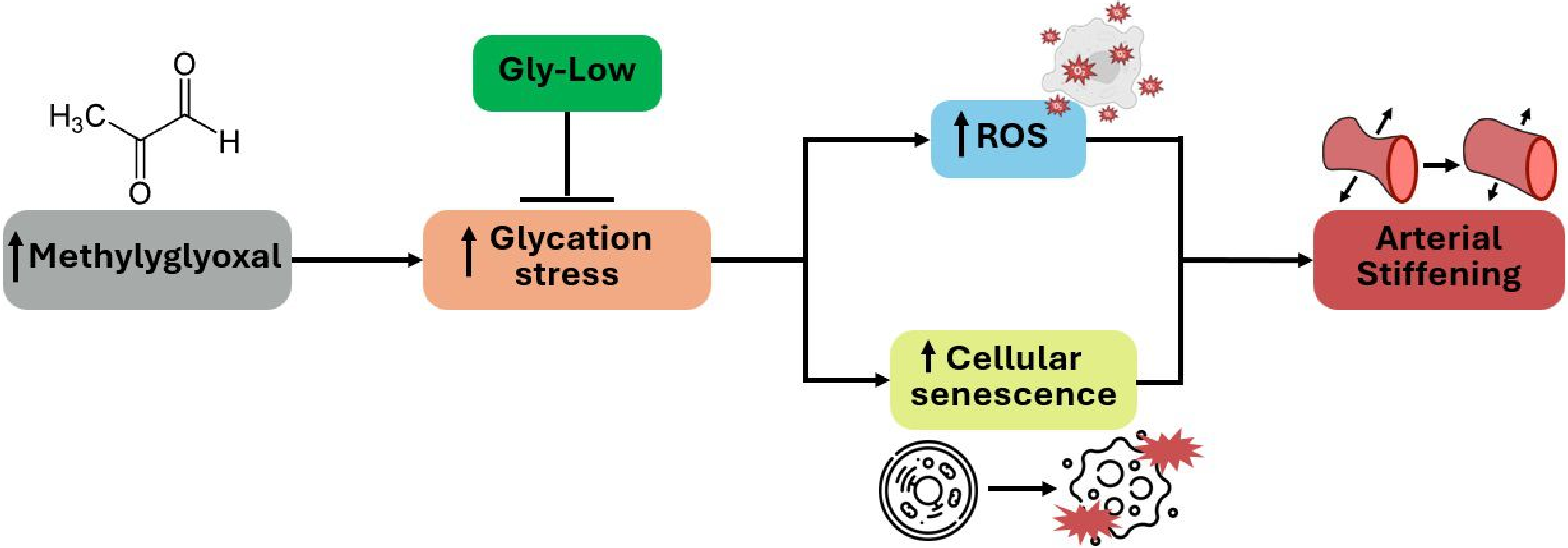
